# Forensics and DNA Barcodes – Do Identification Errors Arise in the Lab or in the Sequence Libraries?

**DOI:** 10.1101/738138

**Authors:** Mikko Pentinsaari, Sujeevan Ratnasingham, Scott E. Miller, Paul D. N. Hebert

## Abstract

Forensic studies often require the determination of biological materials to a species level. As such, DNA-based approaches to identification, particularly DNA barcoding, are attracting increased interest. The capacity of DNA barcodes to assign newly encountered specimens to a species relies upon access to informatics platforms, such as BOLD and GenBank, which host libraries of reference sequences and support the comparison of new sequences to them. As parameterization of these libraries expands, DNA barcoding has the potential to make valuable contributions in diverse forensic contexts. However, a recent publication called for caution after finding that both platforms performed poorly in identifying specimens of 17 common insect species. This study follows up on this concern by asking if the misidentifications reflected problems in the reference libraries or in the query sequences used to test them. Because this reanalysis revealed that missteps in acquiring and analyzing the query sequences were responsible for the misidentifications, a workflow is described to minimize such errors in future investigations. The present study also revealed the limitations imposed by the lack of a polished species-level taxonomy for many groups. In such cases, forensic applications can be strengthened by mapping the geographic distributions of sequence-based species proxies rather than waiting for the maturation of formal taxonomic systems based on morphology.

## Introduction

Species identifications play an important role in forensic analyses in contexts ranging from the interception of trade in CITES-listed species [1] to ascertaining the post mortem interval [2]. There are also expanding opportunities to track the movement of objects and organisms linked to their associated DNA. Although species identifications can play an important role in these contexts, the lack of taxonomic specialists often impedes analysis, a factor which has provoked interest in DNA-based approaches to species identification. Past studies have established that DNA barcodes can often assign specimens to their source species, but have also revealed differences in success among the kingdoms of eukaryotes. For example, the three barcode regions (rbcL, matK, ITS2) for plants deliver lower success than the single gene region (cytochrome *c* oxidase I, COI) used for animals [3]. Because COI generally has high accuracy in species assignment [4–9], the conclusions from a recent study by Meiklejohn et al. [10] were surprising. They assessed the capacity of reference sequences in BOLD, the Barcode of Life Data System [11], and GenBank [12] to generate species-level identifications. Their analysis revealed that both platforms performed similarly in identifying plants and macrofungi, but fared poorly in identifying insect species with BOLD showing lower success than GenBank (35% vs. 53%). By evaluating the factors underpinning the incorrect assignments, the present study revealed that errors in sequence acquisition and interpretation accounted for most, if not all, of the misidentifications. To avoid similar issues in future studies, there is a need to adopt more rigorous procedures for data acquisition and analysis, and to reduce the current reliance on immature taxonomic systems.

## Material and Methods

Meiklejohn et al. [10] analyzed 17 insects including representatives from 12 insect orders – Coleoptera (1), Dermaptera (1), Diptera (5), Ephemeroptera (1), Hymenoptera (1), Lepidoptera (2), Mecoptera (1), Neuroptera (1), Odonata (1), Orthoptera (1), Pthiraptera (1), and Siphonaptera (1). The specimens were obtained from the Smithsonian’s National Museum of Natural History; most were collected 20+ years ago (e.g. *Pediculus humanus* – 1955). Following DNA extraction, the barcode region of COI was PCR amplified and then Sanger sequenced. Reflecting the DNA degradation typical of museum specimens, the sequences recovered were often incomplete (e.g. 254 bp for *Hexagenia limbata*). The resultant sequences were injected into the ID engine on BOLD [11] and into the BLAST function on GenBank [12]. This analysis delivered correct species identifications for six specimens (35%) on BOLD and for nine (53%) on GenBank. The present study was initiated by downloading the 17 sequences from GenBank. They were then resubmitted to the BOLD ID engine and to GenBank BLAST with self matches excluded. Because some of the resultant identifications deviated from those reported in [10], the factors responsible for this discordance were examined.

## Results and Discussion

### ID Results from BOLD and GenBank

Table S1A compares the ID results for the 17 specimens between and those obtained in the present study. The IDs from BLAST matched those reported by [10] as did ten of the IDs from BOLD. The other seven IDs from BOLD corresponded to those from GenBank, but not with the results in [10]. There was a simple explanation for this discordance. Meiklejohn et al. [10] had submitted the reverse complement rather than the coding sequence into the ID engine on BOLD, an approach which generated distant matches. Avoiding this misstep, the number of “correct” identifications generated by BOLD and GenBank was similar (12/17 at the genus level, 9/17 at the species level).

### Factors Responsible for Four ‘Errors’ in Generic Assignment

Both BOLD and GenBank delivered generic identifications deemed incorrect for four specimens. In each case, the query sequence showed close similarity (95–100% in three cases, 90% in one) to taxa belonging to a different order than that analyzed (Supplementary Files 1 & 2). These discordances could either reflect errors in the reference libraries or in the query sequences. The cause for one misidentification was certain; it arose through internal cross-contamination as the sequence for *Hexagenia limbata* was a truncated version of that for *Glossina palpalis* (identical at all 250 bp that overlapped). The other three mismatches involved taxa (springtail, gall midge, strepsipteran) unrepresented among the 17 tested species ruling out internal contamination. Moreover, because of their striking morphological differences to the test taxa (house fly, dragonfly, flea), misidentification can be excluded as a cause. This leaves two possible explanations – contamination in the reference sequence libraries or in the query sequences. Because each query sequence was embedded within many independently generated reference sequences from another order, these cases of misidentification clearly arose from contamination of the query sequences. Cross-contamination is a well-recognized risk when working with museum specimens so it is standard practice to check for its occurrence [13,14], but Meiklejohn et al. [10] make no mention of exercising precautions in this regard. After excluding these four cases, the number of correct identifications for BOLD and GenBank (12/13 for genus, 9/13 for species) was identical.

### Need for Taxonomic Validation of Museum Specimens

The four remaining ‘incorrect’ identifications all involved cases where BOLD and GenBank assigned the query sequence to a species closely related to the taxon analyzed by Meiklejohn et al. [10]. As such, the evidence for misidentification rests on the presumption that their specimens were correctly identified. While the National Museum of Natural History is considered one of the better curated of North American insect collections, the quality of identification of individual specimens depends on the expertise available and the time elapsed since they were assigned to a species. [15]. As such, specimens may be misidentified, mirroring the situation reported in other studies. For example, Meier & Dikow [16] found that 12% of all species-level identifications for a genus of asilid flies from various collections were wrong. Similarly, Muona [17] found that from 1–25% of beetles belonging to two easily discriminated species pairs and one species tetrad were incorrectly identified in a major collection. Similarly, efforts to build a DNA barcode reference library for North American Lepidoptera exposed many misidentified specimens and overlooked cryptic species in major collections [18]. Importantly, all four cases of apparent misidentification reported by Meiklejohn et al. [10] involve species whose recognition is not straightforward. The sole case of generic misidentification involved a presumptive specimen of the cat flea, *Ctenocephalides felis*, whose sequence matched those for the human flea, *Pulex irritans*, on BOLD and GenBank. Because the latter species often uses cats as a host and is morphologically similar to *C. felis*, there is a risk of misidentification. BOLD holds nearly 1,200 records, contributed by 15 institutions, representing four species of *Ctenocephalides* and each possesses a divergent array of barcode sequences. Although the taxonomy of these species is not fully resolved [19], the barcode results support the monophyly of all species in the genus while *P. irritans* forms a sister taxon. Because of the large number of records in the reference library and their derivation from multiple laboratories, the supposed specimen of *C. felis* analyzed by Meiklejohn et al. [10] is almost certainly *P. irritans*. The three remaining cases of presumptive species-level misidentifications involved genera (*Gryllus, Glosssina, Phaenaeus*) with complex taxonomy. One of the three species, *Gryllus assimilans*, was formerly thought to be widely distributed in the New World, but it is now recognized to be a complex of 8+ species, several of which can only be reliably distinguished by their call or life history [20]. Similarly, the query species of tsetse fly (*G. palpalis*) is known to be a complex that includes *G. brevipalpis* [21–24], the species identified by BOLD and GenBank. The third species, *Phanaeus vindex*, is also a complex of at least two species [25], but it is likely more diverse as records for it on BOLD belong to four distinct COI sequence clusters. Because of these taxonomic uncertainties, the four cases of presumptive species- or genus-level misidentifications are best viewed as unconfirmed.

### Resolving Taxonomic Uncertainty

As the preceding section reveals, efforts to assess the resolution of DNA barcodes is often constrained by poor taxonomy. It is certain that some records on BOLD and GenBank derive from misidentified specimens, but there is no easy path to correct them. This fact was powerfully demonstrated by Mutanen et al. [26] study of DNA barcode variation in 4,977 species of European Lepidoptera which revealed that 60% of the cases initially thought to indicate compromised species resolution or DNA barcode sharing actually arose as a result of misidentifications, databasing errors, or flawed taxonomy. As the taxonomic system for European Lepidoptera is very advanced, similar issues will be a greater impediment in most other groups. Databases like BOLD and GenBank record these divergences in taxonomic opinion, but they cannot resolve them, providing strong motivation for approaches that sidestep this barrier. The Barcode Index Number (BIN) system is a good candidate as it makes it possible to objectively register genetically diversified lineages [27]. One of the ‘species’ in the current study, *Forficula auricularia*, provides a good example of the enhanced geographic resolution offered by BINs that could be useful in forensic contexts. This taxon has been known to include two lineages with differing distributions and life histories for >20 years, but it still remains a single recognized species [28,29]. Barcode results indicate that North American populations actually include three divergent lineages with allopatric distributions (Figure 1). As such, BIN assignments provide information on the geographic distributions of the component lineages of this species complex that could be important in certain forensic contexts, but that would be overlooked by a species-based assignment. Because most species of multicellular organisms await description, it is certain that there are many other cases where BIN-based analysis will enhance geographic resolution.

**Figure 1:**
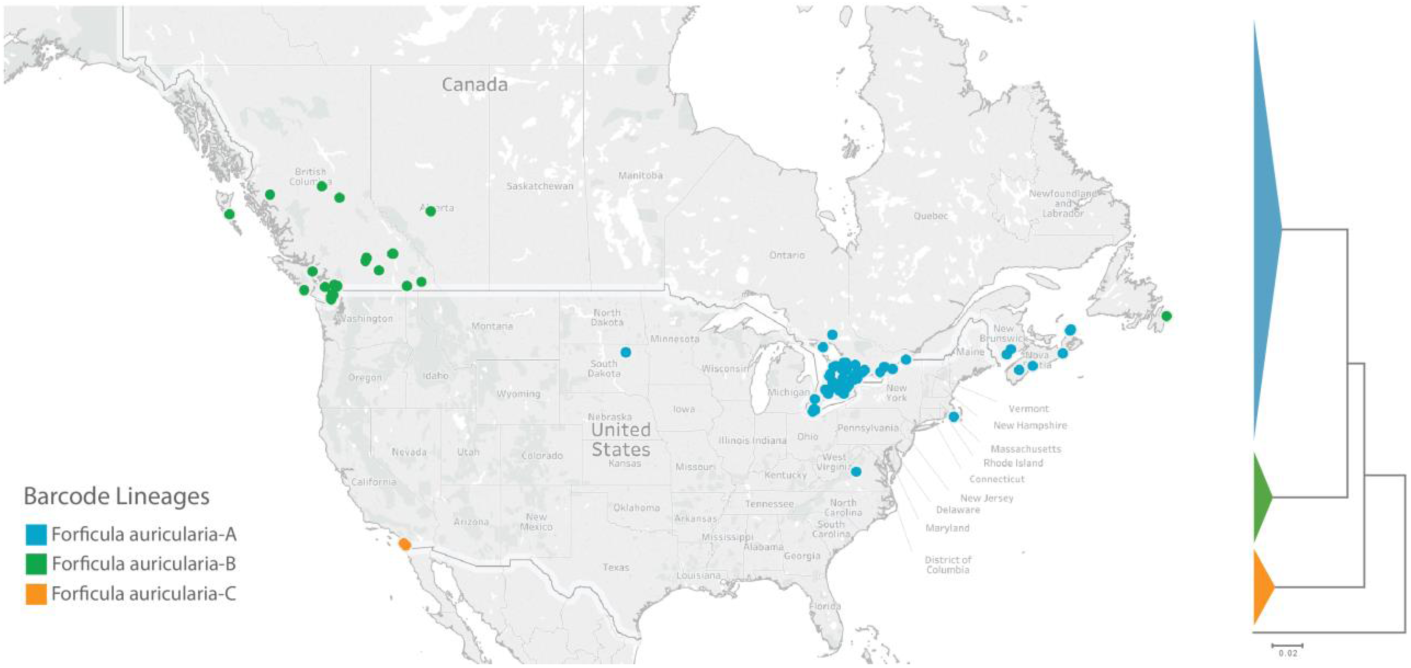
*Geographic distributions and sequence clustering of the three barcode lineages of* Forficula auricularia *in North America*.

### Distinction Between BOLD and GenBank

It is not surprising that BOLD and GenBank demonstrated similar performance in identification, once operational issues were resolved, as many records appear in both platforms. Sequences of COI submitted independently to GenBank are mined and entered into BOLD periodically while records from BOLD are submitted to GenBank when they are published. At present, 11% of all COI barcode records on BOLD originate from GenBank, while 75% of the COI barcodes on GenBank derive from BOLD. Although many records are shared, the two platforms diverge in collateral data. For example, for the 17 species of insects analyzed in [10], 65% of the records originating from BOLD possess GPS coordinates, 60% have trace electropherograms, and 40% have specimen images, while only 26% of those originating from GenBank had GPS coordinates and all lacked images and electropherograms. In addition, BOLD employs BINs to integrate records that lack a genus or species designation with those that possess them. These extended data elements and functionality are a valuable, often essential, component in the evaluation of identification results.

### Conclusions and Path Forward

Six of the 17 species examined by Meiklejohn et al. [10] escaped operational errors, but the other 11 did not (Table 1), explaining the low identification success they reported. Even after correcting for the use of reverse complements, the effectiveness of DNA barcoding could not be evaluated for eight species, those impacted by sequence contamination or taxonomic uncertainty. Importantly, DNA barcode records in BOLD and GenBank did deliver a correct species assignment for the other nine species. While the outcome for these species is reassuring, the lack of an outcome for other taxa reveals the need for improved protocols. Clearly, two conditions need to be satisfied to ensure a correct identification – the query sequences must be legitimate and the reference libraries must be well-validated. As a start, any study that aims to employ DNA barcodes for species identification should include steps to ensure the sequences recovered are valid by including positive and negative controls, by assessing sequence quality, and by checking for contaminants (Figure 2). Presuming the query sequences pass these quality checks, the generation of a reliable identification requires a comprehensive, well-validated reference library. Because BOLD is a workbench for the DNA barcode research community, it will always contain sequences from specimens whose identifications are being refined. The establishment of a Barcode REF library, based upon a small number of carefully validated records for each species, would represent an important step towards improving its capacity to generate reliable identifications. Under ideal circumstances, the reference sequence for each species would derive from its holotype. However, because 90% of all multicellular organisms await description, and the status of many described species groups is uncertain, these efforts will need to be reinforced by a BIN-based approach.

**Table 1:** Three categories of operational errors which compromised efforts by Meiklejohn et al. [8] to test the effectiveness of the BOLD and GenBank reference libraries in identifying 17 insect species.

| Specimen # | ID | Reverse Complement | Contamination | Incorrect ID |
| --- | --- | --- | --- | --- |
| 1 | *Phanaeus vindex* | — | — | Yes |
| 2 | *Forficula auricularia* | Yes | — | — |
| 3 | *Chrysomya rufifacies* | — | — | — |
| 4 | *Calliphora vicina* | — | — | — |
| 5 | *Aedes aegypti* | — | — | — |
| 6 | *Glossina palpalis* | — | — | Yes |
| 7 | *Musca domestica* | Yes | Yes | N.D. |
| 8 | *Hexagenia limbata* | — | Yes | N.D. |
| 9 | *Vespula squamosa* | — | — | — |
| 10 | *Callosamia promethea* | — | — | — |
| 11 | *Danaus plexippus* | — | — | — |
| 12 | *Merope tuber* | Yes | — | — |
| 13 | *Ululodes quadripunctatus* | Yes | — | — |
| 14 | *Gomphus exilis* | Yes | Yes | N.D. |
| 15 | *Gryllus assimilis* | — | — | Yes |
| 16 | *Ctenocephalic felis* | Yes | — | Yes |
| 17 | *Pediculus humanus capitis* | Yes | Yes | N.D. |

**Figure 2:**
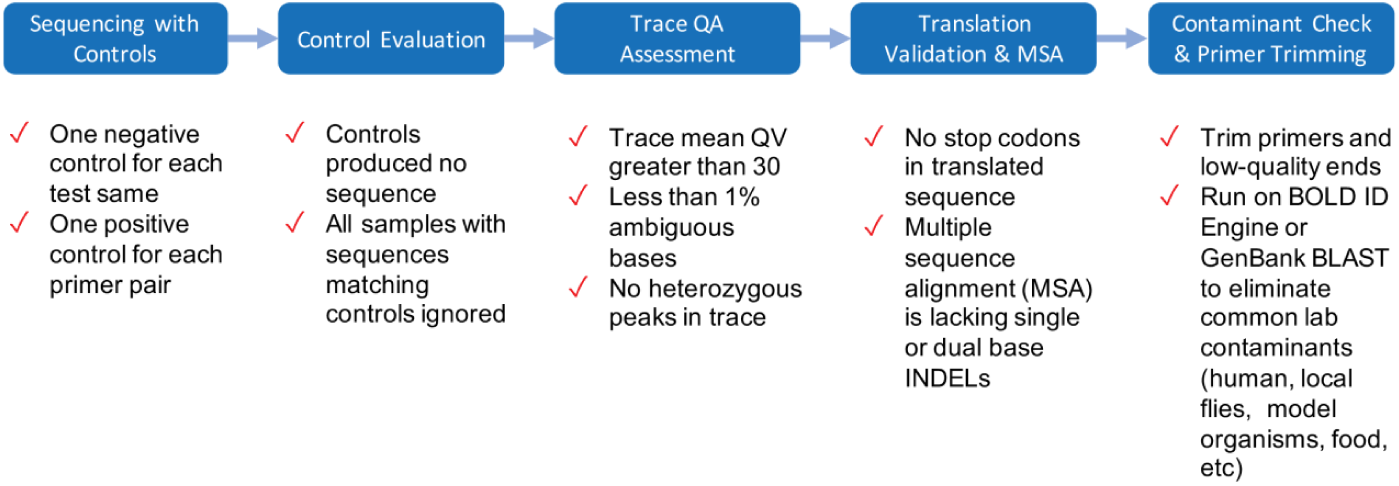
Five key workflow features to maximize the chance of recovering reliable sequence records.

## Acknowledgments

Funding from the Canada First Research Excellence Fund, the Ontario Ministry of Research and Innovation, the Canada Foundation for Innovation, and NSERC support development of the BOLD platform and its computational infrastructure.

## Supplementary Data

Table S1. Comparison of query results (top matches) for 17 insect species between Meiklejohn et al. 2019 (doi:10.1371/journal.pone.0217084) and the present study.

Supplementary file 1 (xlsx). Top 20 matches in GenBank BLAST queries for the four specimens deemed cross-contaminations.

Supplementary file 2 (xlsx). Top 20 matches from queries to the BOLD ID engine for four specimens whose COI sequences derive from cross-contamination

## References

1. Chang C-H, Dai W-Y, Chen T-Y, Lee A-H, Hou H-Y,Liu S-H, et al. DNA barcoding reveals CITES-listed species among Taiwanese government seized chelonian specimens. Genome. 2018 61;615–624.

2. Koroiva R, de Souza MS, Roque FO, Pepinelli M. DNA barcodes for forensically important fly species in Brazil. J Med Entomol. 2018;55: 1055–1061.

3. Coissac E, Hollingsworth PM, Lavergne S, Taberlet P. From barcodes to genomes: extending the concept of DNA barcoding. Mol Ecol. 2016;25: 1423–1428. doi:10.1111/mec.13549

4. Pentinsaari M, Hebert PDN, Mutanen M. Barcoding beetles: A regional survey of 1872 species reveals high identification success and unusually deep interspecific divergences. PLoS One. 2014;9: e108651. doi:10.1371/journal.pone.0108651

5. Hajibabaei M, Janzen DH, Burns JM, Hallwachs W, Hebert PDN. DNA barcodes distinguish species of tropical Lepidoptera. Proc Natl Acad Sci USA. 2006;103: 968–971. doi:10.1073/pnas.0510466103

6. Hausmann A, Godfray HC, Huemer P, Mutanen M, Rougerie R, van Nieukerken EJ, et al. Genetic patterns in European geometrid moths revealed by the Barcode Index Number (BIN) system. PLoS One. 2013;8: e84518. doi:10.1371/journal.pone.0084518

7. Hendrich L, Morinière J, Haszprunar G, Hebert PDN, Hausmann A, Köhler F, et al. A comprehensive DNA barcode database for Central European beetles with a focus on Germany: Adding more than 3,500 identified species to BOLD. Mol Ecol Resour. 2015;15: 795–818. doi:10.1111/1755-0998.12354

8. Huemer P, Mutanen M, Sefc KM, Hebert PDN. Testing DNA barcode performance in 1000 species of European Lepidoptera: large geographic distances have small genetic impacts. PLoS One 2014;9: e115774. doi:10.1371/journal.pone.0115774

9. Kerr KCR, Stoeckle MY, Dove CJ, Weigt LA, Francis CM, Hebert PDN. Comprehensive DNA barcode coverage of North American birds. Mol Ecol Notes. 2007;7: 535–543. doi:10.1111/j.1471-8286.2007.01670.x

10. Meiklejohn KA, Damaso N, Robertson JM. Assessment of BOLD and GenBank – Their accuracy and reliability for the identification of biological materials. PLoS One 2019;14: e0217084. doi:10.1371/journal.pone.0217084

11. Ratnasingham S, Hebert PDN. BOLD: The Barcode of Life Data System (http://www.barcodinglife.org). Mol Ecol Notes. 2007;7: 355–364. doi:10.1111/j.1471-8286.2007.01678.x

12. Benson DA, Cavanaugh M, Clark K, Karsch-Mizrachi I, Lipman DJ, Ostell J, Sayers EW. GenBank. Nucleic Acids Res. 2017 4;45(D1):D37–D42. doi: 10.1093/nar/gkw1070.

13. Siddall ME, Fontanella FM, Watson SC, Kvist S, Erséus C. Barcoding bamboozled by bacteria: Convergence to metazoan mitochondrial primer targets by marine microbes. Syst Biol. 2009;58: 445–451. doi:10.1093/sysbio/syp033

14. Mioduchowska M, Czyz MJ, Goldyn B, Kur J, Sell J. Instances of erroneous DNA barcoding of metazoan invertebrates: Are universal cox1 gene primers too “universal”? PLoS One. 2018;13: e0199609. doi:10.1371/journal.pone.0199609

15. McGinley R J. Where’s the management in collections management? Planning for improved care, greater use and growth of collections. International Symposium and First World Congress on the Preservation and Conservation of Natural History Collections 1993;3: 309–333.

16. Meier R, Dikow T. Significance of specimen databases from taxonomic revisions for estimating and mapping the global species diversity of invertebrates and repatriating reliable specimen data. Conserv Biol. 2004;18: 478–488. doi:10.1111/j.1523-1739.2004.00233.x

17. Muona J. Huomioita eläinmuseon kuoriaskokoelmien virhemäärityksistä [Some observations concerning incorrectly determined beetles in public collections. (Coleoptera)]. Sahlbergia. 2001;6: 34–36.

18. Levesque-Beaudin V, Rosati ME, Silverson N, Warne CP, Brown A, Telfer AC et al. Museum harvesting in major natural history collections. Genome 2017; 60:962. doi: 10.1139/gen-2017-0178

19. Lawrence AL, Brown GK, Peters B, Spielman DS, Morin-Adeline V, Šlapeta J. High phylogenetic diversity of the cat flea (*Ctenocephalides felis*) at two mitochondrial DNA markers. Med Vet Entomol.\ 2014;28: 330–336. doi:10.1111/mve.12051

20. Weissman DB, Gray DA, Pham HT, Tijssen P. Billions and billions sold: Pet-feeder crickets (Orthoptera: Gryllidae), commercial cricket farms, and epizootic densovirus, and government regulations make for a potential disaster. 2012; Zootaxa 3504:67–88.

21. Gooding RH, Krafsur ES. TSETSE GENETICS: Contributions to biology, systematics, and control of Tsetse flies. Annu Rev Entomol. 2005;50: 101–123. doi:10.1146/annurev.ento.50.071803.130443

22. Gooding RH, Solano P, Ravel S. X-chromosome mapping experiments suggest occurrence of cryptic species in the tsetse fly *Glossina palpalis palpalis*. Can J Zool. 2004;82: 1902–1909. doi:10.1139/z05-002

23. Dyer NA, Furtado A, Cano J, Ferreira F, Odete Afonso M, Ndong-Mabale N, et al. Evidence for a discrete evolutionary lineage within Equatorial Guinea suggests that the tsetse fly *Glossina palpalis palpalis* exists as a species complex. Mol Ecol. 2009;18: 3268–3282. doi:10.1111/j.1365-294X.2009.04265.x

24. De Meeûs T, Bouyer J, Ravel S, Solano P. Ecotype evolution in *Glossina palpalis* subspecies, major vectors of Sleeping Sickness. PLoS Negl Trop Dis. 2015;9: e0003497. doi:10.1371/journal.pntd.0003497

25. Price DL. Phylogeny and biogeography of the dung beetle genus *Phanaeus* (Coleoptera: Scarabaeidae). Systematic Entomology. 2009;34: 131–150.

26. Mutanen M, Kivelä SM, Vos RA, Doorenweerd C, Ratnasingham S, Hausmann A, et al. Species-level para- and polyphyly in DNA barcode gene trees: Strong operational bias in European Lepidoptera. Syst Biol. 2016;65(6):1024–1040. doi: 10.1093/sysbio/syw044

27. Ratnasingham S, Hebert PDN. A DNA-based registry for all animal species: the Barcode Index Number (BIN) system. PLoS One. 2013;8:e66213. doi: 10.1371/journal.pone.0066213.

28. Guillet S, Guiller A, Deunff J, Vancassel M. Analysis of a contact zone in the *Forficula auricularia* L. (Dermaptera: Forficulidae) species complex in the Pyrenean Mountains. Heredity. 2000;85: 444–449. doi:10.1046/j.1365-2540.2000.00775.x

29. Guillet S, Josselin N, Vancassel M. Multiple Introductions of the *Forficula auricularia* species complex (Dermaptera: Forficulidae) in eastern North America. Can Entomol. 2000;132: 49–57. doi:10.4039/Ent13249-1

